# RADSex: a computational workflow to study sex determination using Restriction Site-Associated DNA Sequencing data

**DOI:** 10.1101/2020.04.22.054866

**Authors:** Romain Feron, Qiaowei Pan, Ming Wen, Boudjema Imarazene, Elodie Jouanno, Jennifer Anderson, Amaury Herpin, Laurent Journot, Hugues Parrinello, Christophe Klopp, Verena A. Kottler, Alvaro S. Roco, Kang Du, Susanne Kneitz, Mateus Adolfi, Catherine A. Wilson, Braedan McCluskey, Angel Amores, Thomas Desvignes, Frederick W. Goetz, Ato Takanashi, Mari Kawaguchi, H. William Detrich, Marcos Oliveira, Rafael Nobrega, Takashi Sakamoto, Masatoshi Nakamoto, Anna Wargelius, Ørjan Karlsen, Zhongwei Wang, Matthias Stöck, Robert M. Waterhouse, Ingo Braasch, John H. Postlethwait, Manfred Schartl, Yann Guiguen

**Affiliations:** INRAE, LPGP, 35000, Rennes, France; Department of Ecology and Evolution, University of Lausanne, 1015 Lausanne, Switzerland; Swiss Institute of Bioinformatics, 1015 Lausanne, Switzerland; State Key Laboratory of Developmental Biology of Freshwater Fish, College of Life Science, Hunan Normal University, Changsha, China; Department of Systematic Biology, EBC, Uppsala University, Norbyvägen 18D 752 36 Uppsala, Sweden; Institut de Génomique Fonctionnelle, IGF, CNRS, INSERM, Univ. Montpellier, F-34094 Montpellier, France; SIGENAE, Mathématiques et Informatique Appliquées de Toulouse, INRAE, Castanet Tolosan, France; Physiological Chemistry, Biocenter, University of Wuerzburg, 97074 Wuerzburg, Germany; The Xiphophorus Genetic Stock Center, Department of Chemistry and Biochemistry, Texas State University, San Marcos, Texas, USA; Developmental Biochemistry, Biocenter, University of Wuerzburg, 97074 Wuerzburg, Germany; Institute of Neuroscience, University of Oregon, Eugene, OR 97403, USA; Environmental and Fisheries Sciences Division, Northwest Fisheries Science Center, National Marine Fisheries Service, National Oceanic and Atmospheric Administration, 2725 Montlake Blvd East, Seattle, WA 98112, USA; Department of Materials and Life Sciences, Faculty of Science and Technology, Sophia University, 7-1 Kioi-cho, Chiyoda-ku, Tokyo, 102-8554 Japan; Department of Marine and Environmental Sciences, Marine Science Center, Northeastern University, Nahant, Massachusetts 01908; Reproductive and Molecular Biology Group, Department of Structural and Functional Biology, Institute of Biosciences, São Paulo State University, Botucatu, São Paulo, Brazil; Department of Aquatic Marine Biosciences, Tokyo University of Marine Science and Technology, Tokyo 108-8477, Japan; Institute of Marine Research, P.O. Box 1870, Nordnes, NO-5817, Bergen, Norway; Institute of Hydrobiology, Chinese Academy of Sciences, Beijing, China; Leibniz-Institute of Freshwater Ecology and Inland Fisheries, IGB, Berlin, Germany; Department of Integrative Biology, Michigan State University, East Lansing, MI, USA

**Keywords:** sex determination, RAD-Sequencing, fish, computational workflow, visualization

## Abstract

The study of sex determination and sex chromosome organisation in non-model species has long been technically challenging, but new sequencing methodologies are now enabling precise and high-throughput identification of sex-specific genomic sequences. In particular, Restriction Site-Associated DNA Sequencing (RAD-Seq) is being extensively applied to explore sex determination systems in many plant and animal species. However, software designed to specifically search for sex-biased markers using RAD-Seq data is lacking. Here, we present RADSex, a computational analysis workflow designed to study the genetic basis of sex determination using RAD-Seq data. RADSex is simple to use, requires few computational resources, makes no prior assumptions about type of sex-determination system or structure of the sex locus, and offers convenient visualization through a dedicated R package. To demonstrate the functionality of RADSex, we re-analyzed a published dataset of Japanese medaka, *Oryzias latipes*, where we uncovered a previously unknown Y chromosome polymorphism. We then used RADSex to analyze new RAD-Seq datasets from 15 fish species spanning multiple systematic orders. We identified the sex determination system and sex-specific markers in six of these species, five of which had no known sex-markers prior to this study. We show that RADSex greatly facilitates the study of sex determination systems in non-model species and outperforms the commonly used RAD-Seq analysis software STACKS. RADSex in speed, resource usage, ease of application, and visualization options. Furthermore, our analysis of new datasets from 15 species provides new insights on sex determination in fish.

## INTRODUCTION

Sexual reproduction is widespread in animals (Goodenough & Heitman, 2014) and, in gonochoristic species, involves the segregation of the male and female gonadal functions into separate individuals for their entire lives (Bachtrog et al., 2014). Males produce small, mobile gametes while females produce large, immobile gametes, and this difference has fueled the divergent evolution of morphological, physiological, and behavioral traits between the two sexes. Individuals acquire sex-specific traits during sexual development, which starts with sex determination, the process that controls whether an individual develops male or female reproductive organs. Sex determination can be triggered by genetic factors (Genetic Sex Determination, GSD), like in virtually all mammals and birds (Bachtrog et al., 2014), by environmental factors (Environmental Sex Determination, ESD), like temperature in many non-avian reptilia (Pezaro, Doody, & Thompson, 2017), or a combination of the both, as in several fish species (Piferrer, Blázquez, Navarro, & González, 2005). When genetic factors are involved, sex determination is entirely or partially controlled by one or multiple master sex determining (MSD) genes located on sex chromosomes. Genetic sex determination systems can involve male heterogamy (XX/XY and XX/X0), female heterogamy (ZZ/ZW and ZZ/Z0), or multiple loci on different chromosomes (polygenic systems) (Bachtrog et al., 2014). In male- and female-heterogametic systems, the sex chromosomes can be morphologically different, *i.e.* heteromorphic, or similar, *i.e.* homomorphic (Bachtrog et al., 2014). Initially, the overwhelming majority of knowledge on the structure and evolution of sex chromosomes came from studies in mammals (Wallis, Waters, & Graves, 2008) and in *Drosophila* (Salz & Erickson, 2010), where suppression of recombination between the X and Y chromosomes led to degeneration of the Y. These findings have spawned both theoretical and empirical interest in sex chromosomes as models to study the consequences of recombination suppression and associated processes (Bachtrog, 2008; Charlesworth & Charlesworth, 2000; Corcoran et al., 2016; Doorn & Kirkpatrick, 2007; L. Gu, Walters, & Knipple, 2017; Huylmans, Macon, & Vicoso, 2017; Muyle et al., 2012; Peichel et al., 2004). More recent studies investigating sex chromosomes in insects (Blackmon, Ross, & Bachtrog, 2017), non-avian reptilia (Modi & Crews, 2005), amphibians (Miura, 2017), and fishes (Kikuchi & Hamaguchi, 2013) have found many homomorphic sex chromosomes displaying varying levels of differentiation, with the extreme case of a sex locus restricted to allelic variation of a single nucleotide as reported in the Japanese pufferfish (Kamiya et al., 2012). These results called into question the single unified concept of sex chromosome evolution and highlighted the importance of obtaining a broader understanding of sex determination and sex chromosomes in many species across the tree of life. The first step in this process is identifying sex-specific genomic sequences, *i.e.* sequences from the sex locus that are only found in one of the two sexes because of allelic divergence between the sex chromosomes or because of a large insertion in the hemizygous chromosome. These sex-specific sequences can then be aligned to a reference genome to locate the sex locus, identify candidate MSD genes and other genes involved in sex determination, and characterize patterns of differentiation between the sex chromosomes. In addition, facilitating the identification of such sequences has practical applications: sex is an important factor in ecological (Benestan et al., 2017) and conservation studies (Ancona, Dénes, Krüger, Székely, & Beissinger, 2017), as well as agriculture (Al-Ameri, Al-Qurainy, Gaafar, Khan, & Nadeem, 2016; Liao, Yu, & Ming, 2017; Spigler, Lewers, Main, & Ashman, 2008) and animal production (Dan, Mei, Wang, & Gui, 2013; Yano et al., 2013), yet the sex of an individual cannot always be easily determined by non-invasive methods.

Until recently, such analyses were not readily feasible mainly due to technical barriers preventing the precise and high-throughput identification of sex-specific genomic sequences, especially in non-model species, but advances in sequencing technologies have enabled the study of genetic sex determination in a much broader spectrum of taxa by comparing genomes from phenotypically distinct males and females. In essence, this process is akin to comparing genomic differences between two populations or between a mutant and a wild type genotype, and therefore methods from molecular genetics, population genetics, and ecology can be applied. A popular representational approach to comparing the genetics of populations is Restriction Site-Associated DNA Sequencing, or RAD-Seq (Andrews, Good, Miller, Luikart, & Hohenlohe, 2016). RAD-Seq generates short sequences for a small but consistent fraction of the genome and thus allows the sequencing and comparison of multiple individuals from several populations at relatively low cost (Davey et al., 2011) and without requiring additional genomic resources. RAD-Seq has been successfully used to identify sex-specific sequences in non-model species from diverse taxa, including fish (Drinan, Loher, & Hauser, 2018), amphibians (Bewick et al., 2013), non-avian reptilia (Gamble, 2016; Gamble et al., 2017, 2015, 2018; Gamble & Zarkower, 2014; S. V. Nielsen, Banks, Diaz, Trainor, & Gamble, 2018; Stuart V. Nielsen, Daza, Pinto, & Gamble, 2019), invertebrates (Carmichael et al., 2013; Mathers et al., 2015; Pratlong et al., 2017), and plants (Kafkas, Khodaeiaminjan, Güney, & Kafkas, 2015), and to identify the sex chromosomes and the sex locus in some of these species (Wilson et al., 2014). Although dedicated pipelines have been developed to analyze RAD-Seq data specifically for sex determination (Gamble & Zarkower, 2014), most of the above mentioned studies have used STACKS (J. Catchen, Hohenlohe, Bassham, Amores, & Cresko, 2013; J. M. Catchen, Amores, Hohenlohe, Cresko, & Postlethwait, 2011; Hohenlohe, Catchen, & Cresko, 2012; Rochette & Catchen, 2017) to cluster RAD-Seq reads into polymorphic markers subsequently filtered with custom in-house scripts. STACKS, however, requires substantial computational resources to run on large datasets and depends on multiple parameters that can greatly influence the outcome of the analysis (Paris, Stevens, & Catchen, 2017; Rodríguez-Ezpeleta et al., 2016; Shafer et al., 2018), for instance, in the context of sex determination, the classification of a marker as a sex-specific sequence (*e.g.* from a sex-specific insertion) or as a sex-specific allele (*e.g.* an allele specific to one sex in a polymorphic locus) (Utsunomia et al., 2017). In addition, the lack of visualization tools for RAD-Seq data makes the interpretation of results from some datasets difficult, for instance, datasets containing individuals with mis-assigned sex, strong population structure, or sex-bias in sequencing depth.

To overcome these limitations, we developed RADSex, a RAD-Seq data analysis computational workflow specifically designed to study sex determination. RADSex is simple to use, makes no assumptions whether sex-biased markers are sex-biased sequences or sex-biased alleles in a polymorphic locus, requires few resources, and offers visualization tools in the form of an R package. To demonstrate the relevance of RADSex, we analyzed a previously published dataset from the Japanese medaka, *Oryzias latipes*, for which we replicated previous findings, including the identification of two sex-reversed individuals, and we uncovered a previously overlooked Y-specific polymorphism in the sequenced population. We then used RADSex to analyze new datasets that we generated from 15 fish species spanning multiple systematic orders and compared the runtime and memory usage to that of STACKS. In six of these species, we identified the sex-determination system as well as multiple sex markers, and we identified the sex chromosome in one species for which a reference genome was available. Our results show that RADSex is well-suited to study genetic sex-determination in non-model species, outperforming STACKS in speed, resource usage, and ease of use. Furthermore, our multi-species analysis provides insights into the genetic mechanisms of sex-determination in fishes and highlights some of the limitations of RAD-Seq in detecting small sex loci.

## MATERIAL AND METHODS

### Overview of the RADSex analysis workflow

The underlying principle of RADSex is to group identical RAD-Seq reads from all individuals in a dataset into non-polymorphic markers, and then consider the presence or absence of each marker in each individual. Markers created by RADSex thus differ from that of other software such as Stacks (J. M. Catchen *et al.*, 2011) which typically attempt to reconstruct genotypes from polymorphic markers. The RADSex workflow (**Fig. 1**) includes the command-line software *radsex* implemented in C++ (https://github.com/SexGenomicsToolkit/radsex) and the R package *sgtr* (https://github.com/SexGenomicsToolkit/sgtr) to visualize results from *radsex*. The *radsex* software includes several commands, starting with *process*, a data processing command which takes as input a set of demultiplexed reads, *i.e.* one fasta or fastq file containing short reads of consistent size from either single or double digest RAD-seq protocols for each individual. The other commands perform analyses using the output of *process*: *distrib* computes the distribution of marker presence between males and females; *signif* extracts all markers significantly associated with sex; *map* aligns markers to a reference genome sequence; *depth* computes the distribution of marker depths in each individual; *freq* computes the distribution of marker presence in all individuals; and *subset* filters markers based on presence in males and females.

**Figure 1:**
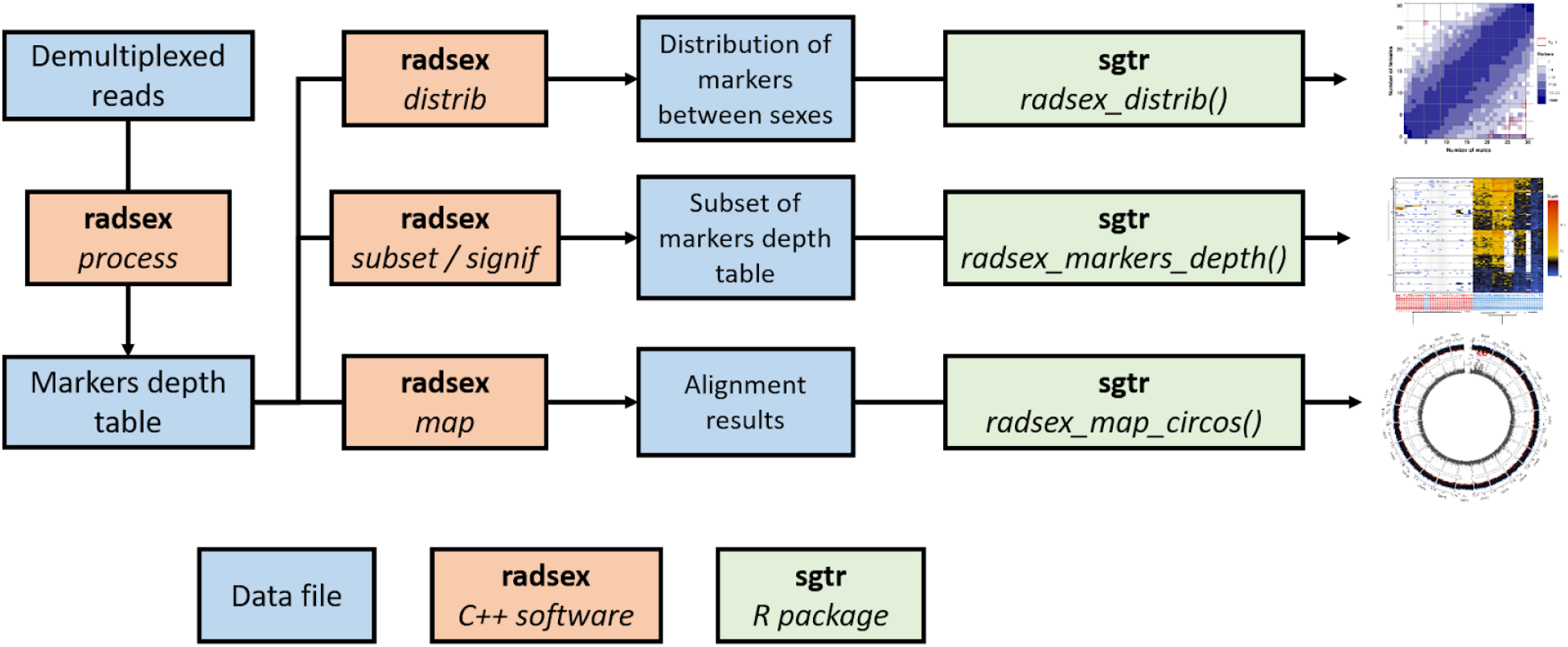
Typical RADSex workflow for sex determination system identification. A table containing the sequencing depth of each marker in each individual is computed using the *process* command from *radsex*. This table is then used as input for the *distrib* command to compute the distribution of markers between males and females, and this distribution is visualized with the *radsex_distrib()* function of *sgtr*. Markers significantly associated with sex are extracted with *signif* and their depth in each individual is visualized with the *radsex_markers_depth()* function of *sgtr.* When a genome assembly is available, markers can be aligned with *radsex map* and results are visualized with the *radsex_map_circos()* function of *sgtr*.

The first step of the workflow is to compute a table of marker depths. The *process* command creates a table with identifier, sequence, and depth in each individual for all unique sequences in the entire dataset. This table summarizes all the sequence information present in the dataset, and each row constitutes a marker for *radsex* (**Supp. Fig. 1**). By design, and in contrast to markers typically generated by other tools like STACKS (J. M. Catchen et al., 2011), markers from *radsex* are not polymorphic. Instead, each allele at a polymorphic locus is treated as a separate marker. The table created by *process* is stored in a plain text tabulated file (**Supplementary table 1**) and is the main input data for all subsequent analyses performed by *radsex*. The *process* command can be parallelized, each thread processing one individual input file at a time.

The next step is to identify markers significantly associated with sex. The *distrib* command uses the table of marker depths generated with *process* to compute the distribution of markers between males and females. Phenotypic sex for each individual is given by a user-supplied tabulated file with the identifier of each individual in the first column and its sex in the second column. In all *radsex* commands, a marker is defined as present in a given individual if the depth of this marker in this individual is higher than a user-specified threshold given by the parameter --min-depth (or -d) (**Supp. Fig. 1**). Following this definition, the *distrib* command enumerates marker presence in every possible combination of number of males and number of females (*e.g.* for a population of one male and two females: 0♂1♀; 0♂2♀; 1♂0♀; 1♂1♀; 1♂2♀), and calculates the probability of association with sex for each combination using Pearson’s chi-squared test of independence with Yates’ correction for continuity. The associated p-value is obtained from the Cumulative Density Function of the chi-squared distribution with one degree of freedom, as implemented in samtools (Li et al., 2009). Bonferroni correction is applied by multiplying the p-value by the total number of markers present in at least one individual. Markers for which association with sex is significant, *i.e. p* < 0.05 (or a user-specified threshold) after Bonferroni correction, can be obtained with the *signif* command and exported either in the same format as the marker depths table or in fasta format. The results of *distrib* (**Supplementary table 2**) can be visualized using the *radsex_distrib()* function from *sgtr*, which generates a tile plot with the number of males on the horizontal axis, the number of females on the vertical axis, and the number of markers represented by a tile’s color for each number of males and females. Tiles for which association with sex is significant are highlighted with a red border. Simplified examples of tile plots for several sex-determining systems are presented in **Fig. 2**.

**Figure 2:**
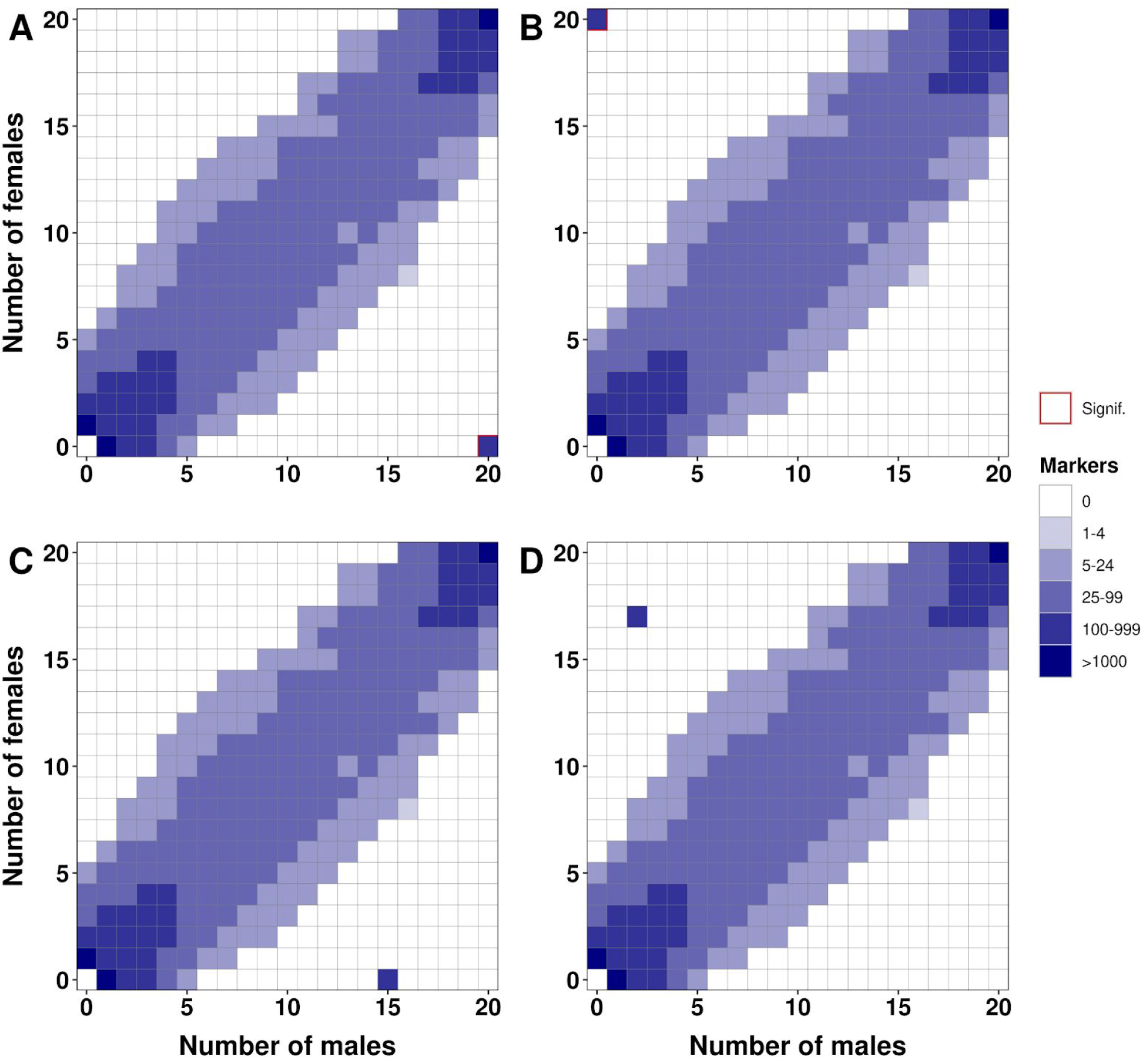
Tile plots showing the theoretical result of *distrib* in four scenarios. Each plot shows the distribution of RADSex markers between phenotypic males (horizontal axis) and phenotypic females (vertical axis), with color intensity for a tile indicating the number of markers present in the corresponding number of males and females. Tiles for which association with phenotypic sex is significant (Chi-squared test, p<0.05 after Bonferroni correction) are highlighted with a red border. **A)** Tile plot for an XX/XY sex determination (SD) system showing markers present in all males and absent from all females in the bottom right corner. **B)** Tile plot for a ZZ/ZW SD system showing markers present in all females and absent from all males in the top left corner. **C)** Tile plot for an XX/XY SD system with five male outliers showing markers present in all but five males and absent from all females at position (15, 0). **D)** Tile plot for a ZZ/ZW SD system with two male and three female outliers showing markers present in all but three females and absent from all but two males at position (2, 17).

Next, RADSex can filter and cluster markers based on depth across individuals to identify consistent absence or presence of markers in some individuals. Markers can be filtered based on presence in males and females using the *radsex subset* command and exported in the same format as the table of marker depths (**Supp. Fig. 1**). After extracting a subset of markers, a heatmap showing the depth of each marker in each individual can be generated using the *radsex_markers_depth()* function from *sgtr*. This function optionally allows the user to cluster both individuals and markers based on depth values and display the resulting cladograms along the heatmap. Distances are computed using the base R *dist* function and clustering is performed with the base R *hclust* function; both distance calculation (*e.g.* euclidean, maximum, binary …) and clustering methods (*e.g.* complete, average, centroid …) can be specified by the user. An application of the *subset* command is the detection of outliers in the sequenced population, *e.g.* sex-reversed individuals or individuals with misassigned sex. In such cases, some markers may have biased presence in individuals from one sex without being statistically associated with phenotypic sex. When clustering individuals based on depth for these sex-biased markers, real outliers will be grouped with individuals from the other sex (see **Fig. 4.B** for an example from a real dataset). Markers obtained with the *signif* command can also be used as input for the *radsex_markers_depth()* function from *sgtr* if they were exported as a table of marker depths.

When a genome assembly is available, markers can be aligned to it using the *radsex map* command in order to locate genomic regions differentiated between sexes. Markers obtained from either the *process*, *signif*, or *subset* commands are aligned to the genome using the *BWA mem* library (Li, 2013), and only markers uniquely aligned with a mapping quality higher than a user-specified threshold are retained. Two metrics are computed for each retained marker: 1) the probability of association with sex p_association with sex’_ estimated from a chi-squared test with Bonferroni correction as described for the *distrib* command, and 2) the sex-bias (*S),* defined as *S* = ♂ / ♂_total_ – ♀ / ♀_total_, where M and F are the number of males and females in which the marker is present, and M_total_ and F_total_ are the total number of males and females in the population, inferred from the user-supplied sex information file. Sex-bias thus ranges from −1 for a marker present in all females and absent from all males to +1 for a marker present in all males and absent from all females, and is zero for a marker present in the same proportion of males and females. Using the output of *radsex map* (**Supplementary table 3**), the *radsex_map_circos()* function of *sgtr* generates a circular plot in which sex-bias, on the top track, and −log_10_(p_association with sex_), on the bottom track, are plotted against genomic position using R’s *circlize* package (Z. Gu, Gu, Eils, Schlesner, & Brors, 2014). Each sector of the plot represents a linkage group, with unplaced scaffolds, which are important to include due to frequent assembly problems with sex chromosomes, appearing in the last sector. Each metric can also be displayed in a Manhattan plot using the *radsex_map_manhattan()* function; in addition, a linear plot showing both metrics for a given genomic region can be generated using the *radsex_map_region()* function from *sgtr*.

Finally, *radsex* provides two commands to assess the distribution of marker depths in the entire dataset. First, the *freq* command computes the number of individuals in which each marker is present for a given minimum depth value and generates a count table of markers present in each number of individuals. Second, the *depth* command computes the minimum, maximum, median, and average marker depth for each individual using markers present in more than 75% of individuals (or a user-specified threshold). Results from *radsex freq* and *radsex depth* can be visualized with the *radsex_freq()* and *radsex_depth()* functions from *sgtr*.

### Sample collection

General information on the different species, species collectors, and samples is given in **Supp. Table 5**.

#### Danios

Several species of the genus *Danio (Danio aesculapii, Danio albolineatus, Danio choprae, Danio kyathit):* were obtained from Eugene Research Aquatics, LLC, Eugene, Oregon, USA. Individuals were euthanized in MS-222 for 10 minutes prior to dissection for sex identification and fin clipping for DNA extraction. Individuals were characterized as male or female based on gonad morphology.

#### Atlantic cod (*Gadus morhua*)

Adult Atlantic cods were sampled from an aquaculture population raised for a nutrition trial (Karlsen et al., 2015). They were sampled at 2.5 years-old in January 2014 after sedation with an overdose of MS222 (200 mg/l) and final euthanasia by applying a sharp blow to the head. Phenotypic sex was determined based on the gonad macroscopy and fin clips were sampled and stored in ethanol until gDNA extraction. Sampling was carried out within the Norwegian animal welfare act guidelines, in accordance with the Animal Welfare Act of 20th December 1974, amended 19th June, 2009, at a facility with permission to conduct experiments on fish (code 93) provided by the Norwegian Animal Research Authority (FDU, www.fdu.no). As this experiment did not involve any treatment apart from euthanasia, no specific permit was required under the Norwegian guidelines.

#### Ayu (*Plecoglossus altivelis*)

Samples were collected in October 2013 in the Jinzu river at Toyama prefecture, Japan. Sexing was performed based on secondary sex characters (male dorsal and anal fins). Sample processing was performed with the approval of the Institutional Animal Care and Use Committee of the Tokyo University of Marine Science and Technology.

#### Spotted gar (*Lepisosteus oculatus*)

Adult spotted gars (total length > 40 cm) were caught by electrofishing in 2014 and 2015 from natural populations around Thibodeaux, Louisiana (Bayou Chevreuil and Atchafalaya basin). Individuals were dissected to confirm sex based on gonadal morphology and fin clips were sampled and stored in 96% ethanol until gDNA extraction. Spotted gar sampling was approved by the University of Oregon Institutional Animal Care and Use Committee (Animal Welfare Assurance Number A-3009-01, IACUC protocol 12-02RA).

#### Pot-bellied seahorse (*Hippocampus abdominalis*)

Adult pot-bellied seahorses were obtained from Seahorse Ways Co. Ltd., Kagoshima, Japan. Individuals were first euthanized in ethyl 3-aminobenzoate methanesulfonate (MS-222) for 5 min before tissue collection. Sex phenotypes were characterized based on gonadal morphology and fin clips were sampled and stored in 90% ethanol until gDNA extraction

#### Common molly (*Poecilia sphenops*)

Individuals were sampled from a laboratory strain (WLC No: 2418) kept by closed colony breeding, established from wild animals collected in the Rio Jamapa. Fish were kept and sampled in accordance with the applicable European Union and National German legislation governing animal experimentation. Experimental protocols were approved through an authorization (568/300-1870/13) of the Veterinary Office. Phenotypic sex was determined by macroscopy of gonads and secondary sex characters, including male pigmentation (black spotting, androgen dependent pattern, not expressed in females), male gonopodium (sex-specific anal fin morphological differentiation), and female gravidity spot.

#### Black widow tetra (*Gymnocorymbus ternetzi*)

Individuals were collected in the Guaporé River (Pontes e Lacerda, Mato Grosso, Brazil) in December 2015. Fish were caught using trawl nets (3 mm mesh size, 25 × 2.5 m) which were placed at two points of the river. Samples were then euthanized with benzocaine hydrochloride solution of > 250 mg/L and preserved into glasses containing 100% ethanol for later gDNA extraction and histological analysis. Phenotypic sex was determined from histological processing of the gonads following standard protocols: the material was dehydrated in a series of alcohol, embedded in Leica historesin (methacrylate glycol) (Leica Microsystems, Wetzlar, Germany), sectioned with 3μm thickness, and stained with Hematoxylin/Eosin for examination under microscope. All procedures were consistent with Brazilian legislation regulated by the National Council for the Control of Animal Experimental (CONCEA) and Ethical Principles in Animal Research (Protocol n. 671-CEUA).

#### Banded knifefish (*Gymnotus carapo*)

Samples were collected along the Piracicaba River (Piracicaba, São Paulo, Brazil) in December 2014. The specimens were collected using sieves with square (2 × 1 m) and round (1.2 m diameter) shape meshes. Fish were then transported to the Department of Structural and Functional Biology, Institute of Biosciences, Botucatu (São Paulo, Brazil). Samples were euthanized with benzocaine hydrochloride solution of > 250 mg/L and preserved into glasses containing 100% ethanol for later gDNA extraction and histological analysis. Phenotypic sex was determined from histological processing of the gonads following standard protocols: the material was dehydrated in a series of alcohol, embedded in Leica historesin (methacrylate glycol) (Leica Microsystems, Wetzlar, Germany), sectioned with 3 μm thickness, and stained with Hematoxylin/Eosin for examination under microscope. All procedures were consistent with Brazilian legislation regulated by the National Council for the Control of Animal Experimental (CONCEA) and Ethical Principles in Animal Research (Protocol n. 671-CEUA).

#### Walleye (*Sander vitreus*)

Fin clips were obtained from individuals from the Spirit Lake strain collected by gillnet in April 2015 from Clear Lake, Iowa (USA). These fish were part of an annual broodstock collection to obtain gametes for juvenile production used in stocking inland lakes. The sex of the fish was determined by a combination of the presence of expelled semen or ovulated eggs, dissection for fish that were dead, and size for a small number of fish.

#### Marbled notothen (*Notothenia rossii*)

Reproductively active individuals were collected in April-May 2013 and 2014 by bottom trawling and baited traps deployed from the ARSV *Laurence M. Gould* in the vicinity of Low Island and Dallmann Bay on the West Antarctic Peninsula. Live fish were transferred immediately from the trawl net or trap to the ship aquaria and transported to the aquatic facilities at Palmer Station, Antarctica, where they were maintained in flow-through seawater tanks at ambient temperatures of −1 to +1 °C. Following euthanasia using MS-222 (Syndel, Ferndale, WA, USA), individuals were dissected to identify the sex by gonad examination and a fragment of the gonads was preserved in Bouin’s fixative for histological verification of the sex at the University of Oregon. A fin clip was also sampled and stored in 80% ethanol for gDNA extraction. All procedures were performed according to protocols approved by the Institutional Animal Care and Use Committees (IACUC) of the University of Oregon (#13-27RRAA).

#### Tench (*Tinca tinca*) and common carp (*Cyprinus carpio*)

Tenchs (*Tinca tinca*) were sampled in January 2017 from an aquaculture stock from the fish production unit of the fishing federation of Ille-et-Vilaine (Pisciculture du Boulet, Feins, France). A local aquaculture strain of common carp, *Cyprinus carpio* (U3E-INRA experimental aquaculture strain) reared in outdoor ponds was sampled in November and December 2013. For both species, phenotypic sex was recorded on each animal either by the presence of expelled semen or eggs or by macroscopic examination of the gonad after euthanasia with a lethal dose of MS-222 and dissection of the fish. Fin clips were taken on each animal and stored in ethanol 90% until gDNA extraction. For both common carp and tench, sampling and euthanasia procedures conformed to the principles for the use and care of laboratory animals, in compliance with French (“National Council for Animal Experimentation” of the French Ministry of Higher Education and Research and the Ministry of Food, Agriculture, and Forest) and European (European Communities Council Directive 2010/63/UE) guidelines on animal welfare. No specific permit was required under French guidelines for a simple euthanasia and tissue sampling.

### Genomic DNA (gDNA) extraction

Genomic DNA (gDNA) of *Lepisosteus oculatus*, *Plecoglossus altivelis*, *Gadus morhua*, *Notothenia rossii*, *Sander vitreus*, *Tinca tinca*, *Carassius auratus*, and all *Danios* were extracted from fin clips stored in ethanol using NucleoSpin Kits for Tissue (Macherey-Nagel, Duren, Germany) following the producer’s protocol. Genomic DNA of *Gymnotus carapo*, *Gymnocorymbus ternetzi*, *Hippocampus abdominalis*, and *Poecilia sphenops* were extracted from individual fin clips stored in ethanol or from pooled organs of individual fish for *P. sphenops* with a phenol/chloroform protocol. Fin clips or tissues were lysed in 1 ml extraction buffer (0.1 M EDTA pH 8, 0.2% SDS, 0.2 M NaCl) containing 200 μg/ml proteinase K for three hours at 80°C, and 500 μL phenol was added to each sample. After mixing and incubating at room temperature for 10 min, 500 μL chloroform/isoamyl alcohol (24:1) was added. Samples were incubated with occasional inversion at room temperature for 10 min and then centrifuged for 10 min at 5000 g (5°C). The upper layer was transferred to a new tube and one volume of chloroform/isoamyl alcohol (24:1) was added. The samples were kept at room temperature for 10 min with occasional inverting, and subsequently centrifuged for 10 min at 5000 g (5°C). The supernatant was transferred to a glass vial on ice and 2.5 times the sample volume of cold 100% ethanol was carefully added. The DNA was spooled with a glass rod and dissolved in TE buffer pH 8. All gDNA concentrations were quantified with a NanoDrop ND2000 spectrophotometer (Thermo scientific, Wilmington, Delaware) and/or Qubit (ThermoFisher, France) before processing gDNA for RAD libraries construction.

### RAD-Sequencing and demultiplexing of RAD-Seq reads

RAD libraries were constructed for each species from individual fish gDNA using a single restriction enzyme, SbfI, following the standard protocol as previously described (Amores, Catchen, Ferrara, Fontenot, & Postlethwait, 2011). Libraries were sequenced as single end 100 bp reads on one lane of Illumina HiSeq 2500. Quality control and demultiplexing of the reads was performed with the *process_radtags.pl* wrapper script from Stacks version 1.44 (J. M. Catchen et al., 2011) using the following options: discard reads with low quality scores (-q), remove any reads with an uncalled base (-c), and rescue barcodes and RAD-seq reads (-r). Demultiplexing results for each species are summarized in **Supplementary Table 7**.

### Performance measurements

Runtime and peak memory usage of *radsex* (version 1.0.0) and Stacks (version 2.2) were measured with “/usr/bin/time -v <command>” on the Genotoul computational platform (Toulouse, France) using 16 threads. For *radsex*, performance was measured for the *process*, *distrib*, and *signif* commands run sequentially with a minimum depth of 1, as this value would require the longest runtime and memory. For Stacks, performance was measured for the *denovo_map.pl* pipeline with the following settings: “-M 3 -n 2 -X populations:--fstats”, which represent a standard usage to identify sex-specific markers.

### Software used in the analyses

All analyses were performed using version 1.0.0 of *radsex* and figures were generated using version 1.0.1 of the *sgtr* R package in R version 3.5.2. Both *radsex* and *sgtr* are released under GPLv3 license; releases and source code are available at https://github.com/SexGenomicsToolkit/radsex for *radsex* and https://github.com/SexGenomicsToolkit/sgtr for *sgtr*. Both *radsex* and *sgtr* are also available on the conda channel Bioconda. A complete documentation for RADSex is available at https://sexgenomicstoolkit.github.io/html/radsex/introduction.html.

## RESULTS

### Performance comparison of RADSex and Stacks

The runtime and peak memory usage of *radsex process, distrib,* and *signif* were measured on datasets from 15 fish species that we generated for this study and compared to that of STACKS *denovo_map.pl* (version 2.2, (J. Catchen et al., 2013)). Using our configuration, RADSex’s runtime was linearly and positively correlated to the number of RAD-seq reads, ranging from 139 seconds for *Tinca tinca*, whose dataset contained ~ 130 M. RAD-seq reads, to 1106 seconds (18 min and 26 s) for *Sander vitreus*, whose dataset contained ~ 262 M. RAD-seq reads (**Fig. 3.A**). RADSex was on average 42 times faster than STACKS on the same dataset, ranging from 26 times faster for *Danio choprae* to 88 times faster for *Danio albolineatus* (**Fig 3.B**). RADSex’s peak memory usage was also linearly and positively correlated to the number of RAD-seq reads, ranging from 2.4 Gb for *Poecilia sphenops* to 17 Gb for *S. vitreus* (**Fig 3.C**). RADSex used on average 1.9 times less memory than STACKS to analyze the same dataset, ranging from 1.2 times less memory for *D. choprae* to 3.4 times less memory for *D. albolineatus* (**Fig 3.D**). It is worth noting that for *radsex,* the peak memory usage and most of the runtime were in the *process* command, which needs to be run only once for each dataset because filtering parameters (*i.e.* minimum depth for marker presence) can be applied in the later steps of the RADSex pipeline.

**Figure 3:**
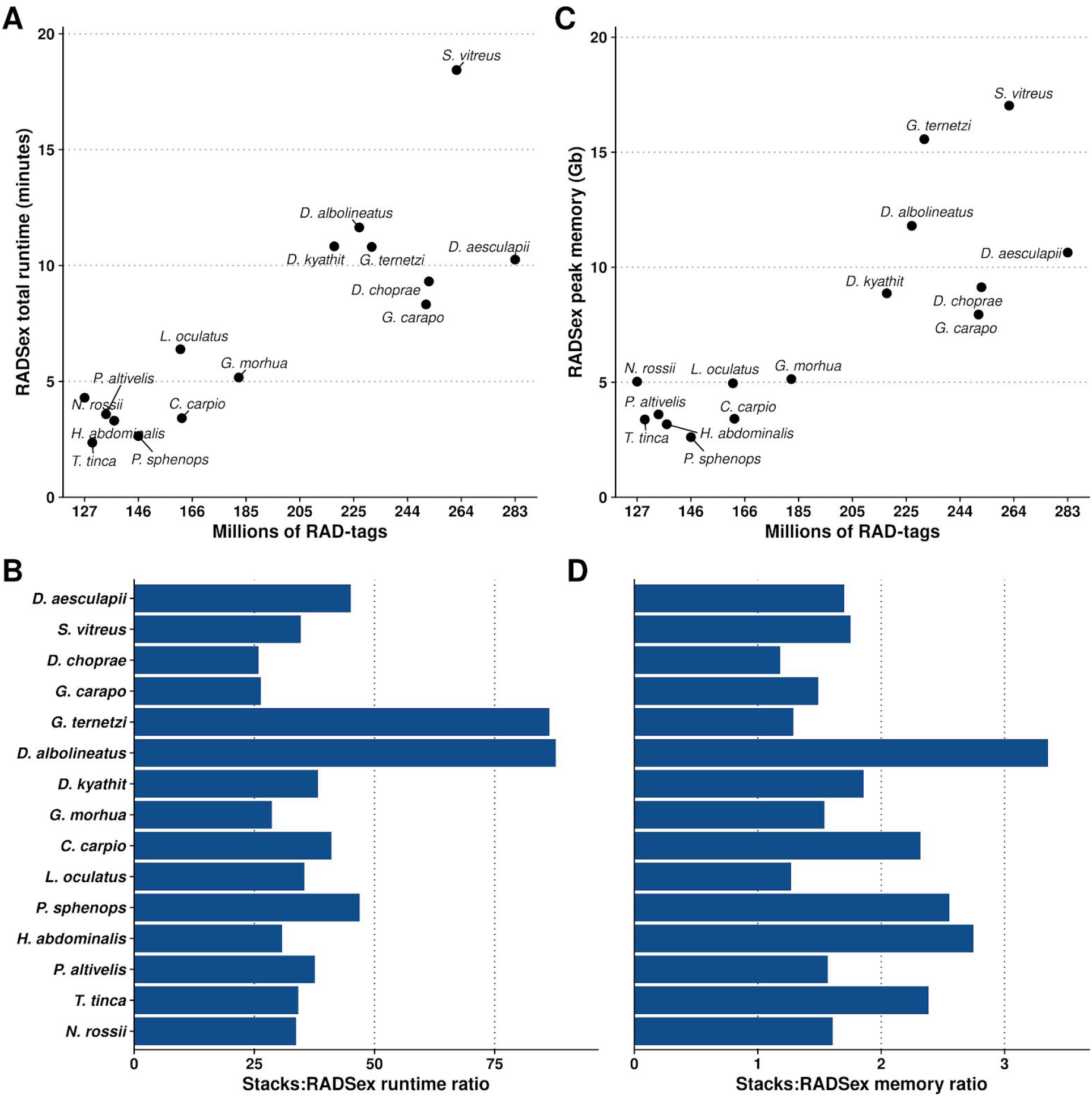
Performance of the RADSex workflow (*process, signif,* and *distrib*) and comparison with *Stacks*. **A**: Total runtime of RADSex (in minutes) plotted against the number of reads for each dataset. **B**: Peak memory of RADSex in gigabytes plotted against the number of RAD-seq reads for each dataset **C**: Runtime ratio between *Stacks* (version 2.2) and RADSex showing that RADSex was 25 to 80 times faster than *Stacks*. Datasets are sorted by number of RAD-seq reads. **D**: Peak memory ratio between *Stacks* and RADSex showing that RADSex used 1.2 to 3.4 times less memory than *Stacks*. Datasets are sorted by number of RAD-seq reads.

### RADSex validation on a published dataset

To assess whether RADSex can recover previous results on sex determination, we analyzed a dataset which has been used to characterize a major sex-determination locus with a STACKS-based approach (Wilson et al., 2014). This dataset consists of RAD-Seq reads from 31 males and 30 females from the *carbio* strain of *Oryzias latipes*, the Japanese medaka, which has a known XX/XY sex determination system with chromosome 1 as the sex chromosome (Matsuda et al., 2002; Indrajit Nanda et al., 2002). We analyzed this *O. latipes* dataset with RADSex using a minimum depth of 10 (-d 10) to define the presence of a marker in an individual. This value of minimum depth was chosen based on the median sequencing depth in each individual computed with *radsex depth*: the average median sequencing depth was 34, and thus a stringent minimum depth of 10 can be used to discard markers from potential sequencing errors but still retain real markers. In total, we found 121,492 markers present in at least one individual with a minimum depth of 10, among which 194 markers were significantly associated with male phenotype (*radsex signif*, p < 0.05, chi-squared test with Bonferroni correction, highlighted tiles in **Fig 4.A**). Among these 194 markers, 165 were present in at least 20 of the 31 males. In addition, a single marker present in 26 of the 30 females and 5 males was significantly associated with female phenotype (**Fig. 4.A**). However, we did not find any marker simultaneously present in more than 29 of the 31 phenotypic males and absent from all phenotypic females. To understand why no marker was found exclusively in all males given that medaka has an XY/XX sex determination system, we extracted individual depth for all 165 markers present in at least 20 males and none of the females using *radsex subset*. We clustered both markers and individuals separately based on these depth values and generated a heatmap of the results using the *radsex_markers_depth()* function of *sgtr* (**Fig. 4.B**). As expected, phenotypic males and females clustered in two separate groups, except for two phenotypic males (green arrows in **Fig. 4.B**) that clustered with the phenotypic females, showing zero or very low depth for the extracted markers. These two outlier phenotypic males had already been found to be sex-reversed genetic XX females by retrospective genotyping in Wilson *et al.* (Wilson et al., 2014). After correcting the designated phenotypic sex of these two individuals, *radsex signif* yielded 232 markers significantly associated with male phenotype and no marker significantly associated with female phenotype (p < 0.05, chi-squared test with Bonferroni correction), as expected for an XY/XX system. In addition, six males forming a sub-cluster within the male cluster from *sgtr radsex_markers_depth()* had low or null depth for almost half of the extracted markers, suggesting that several Y chromosome haplotypes could be present in the sequenced population.

**Figure 4:**
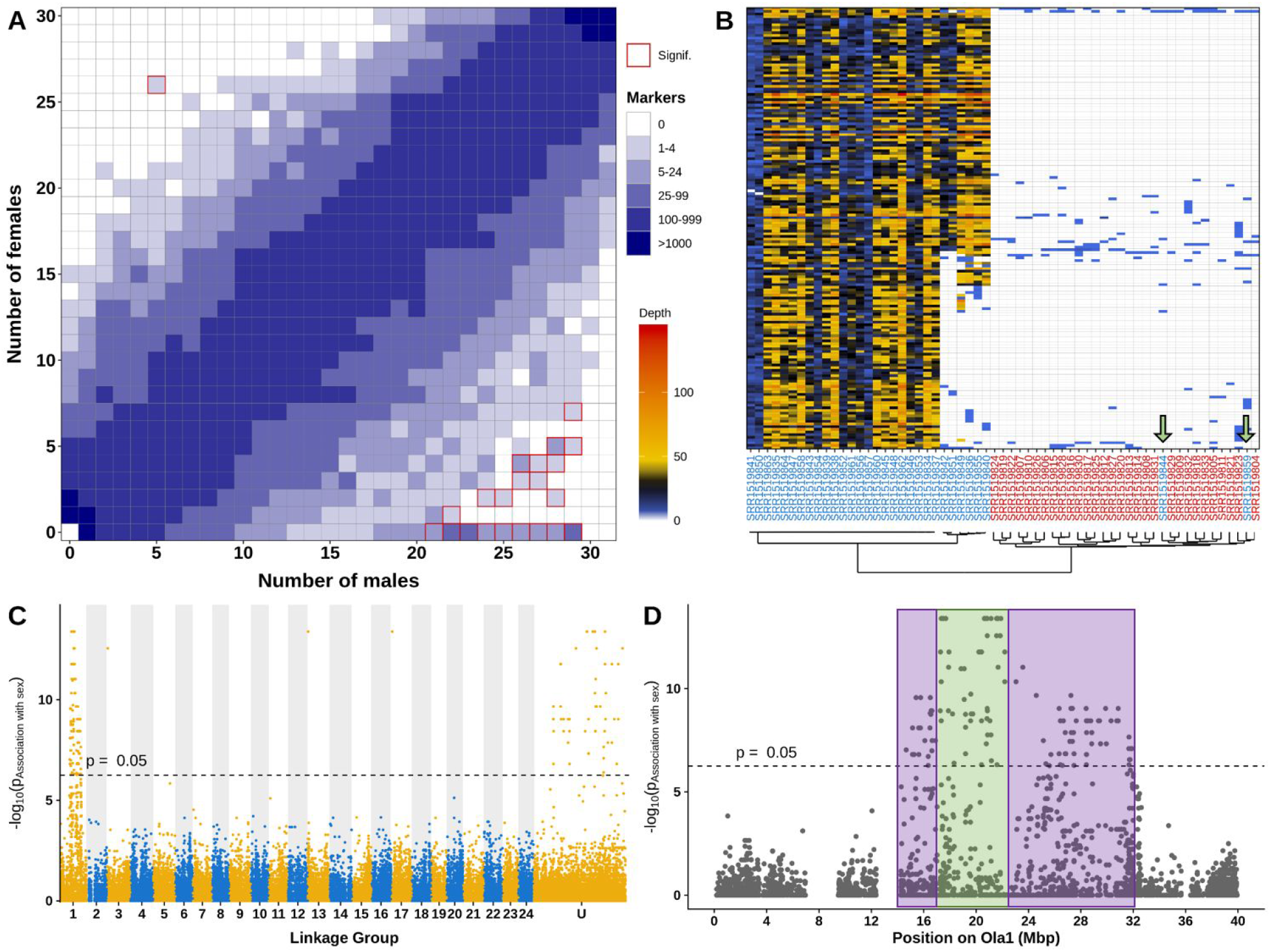
Validation of the RADSex approach on the *Oryzias latipes* dataset. **A**: Tile plot showing the distribution of RADSex markers between males (horizontal axis) and females (vertical axis) using the original sex assignment and a minimum depth of m = 10. Color intensity indicates the number of markers present in the corresponding number of males and females, and a red border indicates significant association with sex (Chi-squared test, p<0.05 after Bonferroni correction). **B**: Heatmap showing individual depth for all markers present in at least 20 males and absent (-d 10) from all females. Each row corresponds to a marker, and each column corresponds to an individual. Males (blue labels) and females (red labels) clustered in two separate groups, except for two male outliers, identified by the green arrows, which clustered with the females. In addition, six males forming a sub-cluster within the male cluster had low or null depth for half of the markers. **C**: Manhattan plot showing −log10(P) of a chi-squared test on number of males and number of females, after Bonferroni correction, for all RADSex markers aligned to the genome of *O. latipes*, with odd linkage groups (LGs) in yellow with light background and even LGs in blue with shaded background. Unplaced scaffolds were combined in a super scaffold (U) by order of decreasing size. The dashed line indicates a p-value of 0.05 after Bonferroni correction. Most markers significantly associated with sex were aligned to LG01 (Ola1) or to unplaced scaffolds. **D**: The −log10(P) of a chi-squared test on the number of males and number of females is plotted against genomic position on LG01 of *O. latipes* (Ola1) for all RADSex markers aligned to this LG. The dashed line indicates a p-value of 0.05 after Bonferroni correction. Markers significantly associated with sex were aligned to a continuous region between 14.7 and 31.9 Mb. Markers present in all males apart from the two outliers and absent in all females were aligned in a continuous region spanning from 17 to 23 Mb (green box), while male-biased markers absent from the sub-cluster of six males were aligned in two regions, from 14 to 17Mb and from 24 to 31.9 Mb (purple boxes).

To identify the sex chromosome and to delimit the sex-differentiated region, we aligned the RADSex markers to the assembly used by Wilson *et al.* (MEDAKA1, available at http://77.235.253.122:8012/Oryzias_latipes/Info/Index) using *radsex map.* Among the 232 markers significantly associated with male phenotype after reassignment of the sex-reversed fish, 131 (56%) aligned to LG01 of *O. latipes* (Ola1), three (1%) aligned to other chromosomes, 54 (23%) aligned to unplaced scaffolds, and 44 (19%) were not uniquely aligned with mapping quality higher than 20 (**Fig. 4.C**). On Ola1, markers significantly associated with male phenotype aligned to a continuous region spanning from 14.7 Mb to 31.9 Mb (**Fig. 4.D**). Within this region, markers found in all males and no females aligned between 17 and 23 Mb (green box in **Fig. 4.D**) and markers present in all but the six males identified in the clustering step and absent from all females aligned between 14.7 Mb and 31.9 Mb (purple boxes in **Fig. 4.D**). This result supports the hypothesis that multiple Y chromosome haplotypes are present in this medaka population, with one region located between 17 and 23 Mb on Ola1 showing strong X/Y differentiation in all Y haplotypes a wider region located between 14.7 and 31.9 Mb on Ola1 showing strong X/Y differentiation in only one Y haplotype.

Overall, these results are consistent with Wilson *et al.* (Wilson et al., 2014), who found 248 RAD-seq reads associated with male phenotype mapping to a region ranging from 14.3 to 32.5 Mb on Ola1. However, our analysis revealed a previously unidentified Y-specific polymorphism in the sequenced population. This example highlights the effectiveness and versatility of RADSex and its visualization tools to identify, explore, and explain patterns in datasets that can be overlooked with traditional approaches.

### Analysis of 15 new ray-finned fish RAD-sequencing datasets

RAD-Seq datasets were generated for 15 ray-finned species in which no markers for genetic sex determination were available when we initiated this work. For each species, we used RADSex to search for sex-biased markers with a minimum marker depth (-d) of 1, 2, 5, and 10. We report results for a minimum marker depth of 10 (**Table 1**) as it is the most stringent, except in three species with the lowest sequencing depths, for which we report results for a minimum marker depth of 2 (**Supp Fig. 2**). We identified an XX/XY sex determining system in six species (*Cyprinus carpio*, *Gymnotus carapo*, *Plecoglossus atlivelis*, *Tinca tinca*, *Gadus morhua*, and *Poecilia sphenops*) with a few male and/or female outliers in four of these species (**Fig. 5**). Markers significantly associated with phenotypic sex for each species are provided in **Supplementary table 6**. An XX/XY SD system is in agreement with previous reports for *C. carpio* (Gomelsky, 2003), *G. carapo* (da Silva, Matoso, Artoni, & Feldberg, 2014), *P. altivelis* (Watanabe, Yamasaki, Seki, & Taniguchi, 2004), and *G. morhua* (Haugen et al., 2012; Whitehead, Benfey, & Martin-Robichaud, 2012), but conflicts with the identification of *P. sphenops* being female heterogametic (I. Nanda, Schartl, Epplen, Feichtinger, & Schmid, 1993). However, opposite population-specific sex determination systems have also been described in *P. sphenops* (Volff & Schartl, 2001), and because previous studies were performed on ornamental fish and laboratory strains, previously derived conclusions may not apply to wild populations like the one we analyzed. Lastly, to the best of our knowledge, sex determination was not previously characterized in the tench, *Tinca tinca*.

**Table 1:** Summary of RADSex results for the 15 datasets analyzed. Species where a sex-determination system was identified with *radsex* are highlighted with a grey background. The number of markers significantly associated with phenotypic sex is also given for each species, with the value of ‘*d*’ indicating to the minimal number of reads for a marker to be considered present in an individual in the analysis.

| Species | SD system identified with *radsex* | Number of sex markers (d=) |
| --- | --- | --- |
| ***Cyprinus carpio*** | **XX/XY with male outliers** | **7 (d = 10)** |
| *Danio aesculapii* | No SD identified | 0 (d = 10) |
| *Danio albolineatus* | No SD identified | 0 (d = 10) |
| *Danio choprae* | No SD identified | 0 (d = 10) |
| *Danio kyathit* | No SD identified | 0 (d = 10) |
| ***Gadus morhua*** | **XX/XY with outliers** | **3 (d = 10)** |
| *Gymnocorymbus ternetzi* | No SD identified | 0 (d = 2) |
| ***Gymnotus carapo*** | **XX/XY with female outliers** | **8 (d = 10)** |
| *Hippocampus abdominalis* | No SD identified | 0 (d = 10) |
| *Lepisosteus oculatus* | No SD identified | 0 (d = 2) |
| *Notothenia rossii* | No SD identified | 0 (d = 10) |
| ***Plecoglossus altivelis*** | **XX/XY** | **47 (d = 10)** |
| ***Poecilia sphenops*** | **XX/XY with female outliers** | **7 (d = 10)** |
| *Sander vitreus* | No SD identified | 0 (d = 2) |
| ***Tinca tinca*** | **XX/XY** | **6 (d = 10)** |

**Figure 5:**
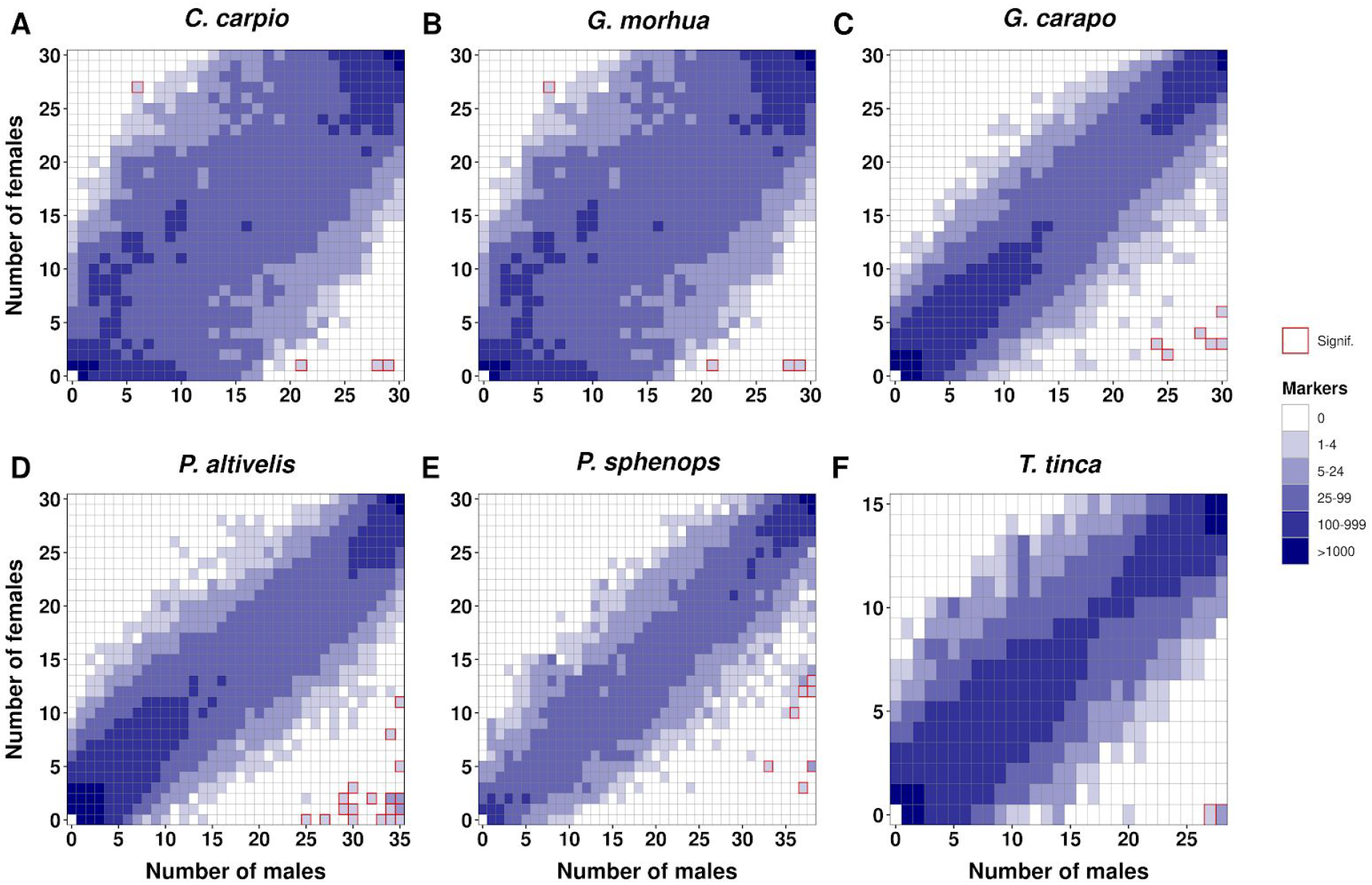
Distribution of markers between males and females for six species (A-F) with markers significantly associated with sex. The number of markers, represented by tile color intensity, is plotted for every combination of number of males (horizontal axis) and number of females (vertical axis) for each species. Combinations of number of males and number of females for which association with sex is significant (p<0.05, Chi-squared test with Bonferroni correction) are highlighted in red. Distributions were computed with a minimum depth of 10 (-d 10).

In *G. morhua*, three markers were significantly associated with male phenotype, but they were also found in two females and none of them was present in more than 27 of the 34 males **(Fig. 5.B)**. Furthermore, multiple markers were found predominantly in males but not significantly associated with sex, with eight markers consistently present with depth higher than five in 20 males and absent from all females (these markers are highlighted with arrows in **Fig. 6.A**). The three markers significantly associated with sex (red-outlined boxes) and most other markers strongly but not significantly associated with sex aligned to cod linkage group 11 (Gmo11) (**Fig. 6.B and 6.C**). Markers strongly associated with sex aligned to a region spanning from 0 to 20 Mb on Gmo11, and the three markers significantly associated with sex aligned between 10 to 15 Mb on this chromosome, in a region that was previously characterized as the region containing the sex locus in this species (Kirubakaran et al., 2019; Star et al., 2016). These results confirm the previously identified XX/XY sex determining system and Gmo11 as the sex chromosome in Atlantic cod (Haugen et al., 2012; Whitehead et al., 2012), but our findings suggest an additional sex determination complexity in the studied Atlantic cod aquaculture population which had not been observed before on wild populations of Atlantic cod (Kirubakaran et al., 2019; Star et al., 2016). The incomplete sex-linkage of sex-biased markers may be explained by a Y chromosome polymorphism and/or the existence of sex-reversed genetic XX females. Although Y chromosome population differences have rarely been explored, at least in fish species, the results from our re-analysis of a medaka population suggest that such polymorphism may be more frequent than expected. In addition, female-to-male sex-reversal could be a consequence of intensive rearing aquaculture conditions leading to stress induced sex-reversal (Geffroy & Douhard, 2019), which has been observed in other fish species reared in laboratory or aquaculture facilities (Pan et al., 2019).

**Figure 6:**
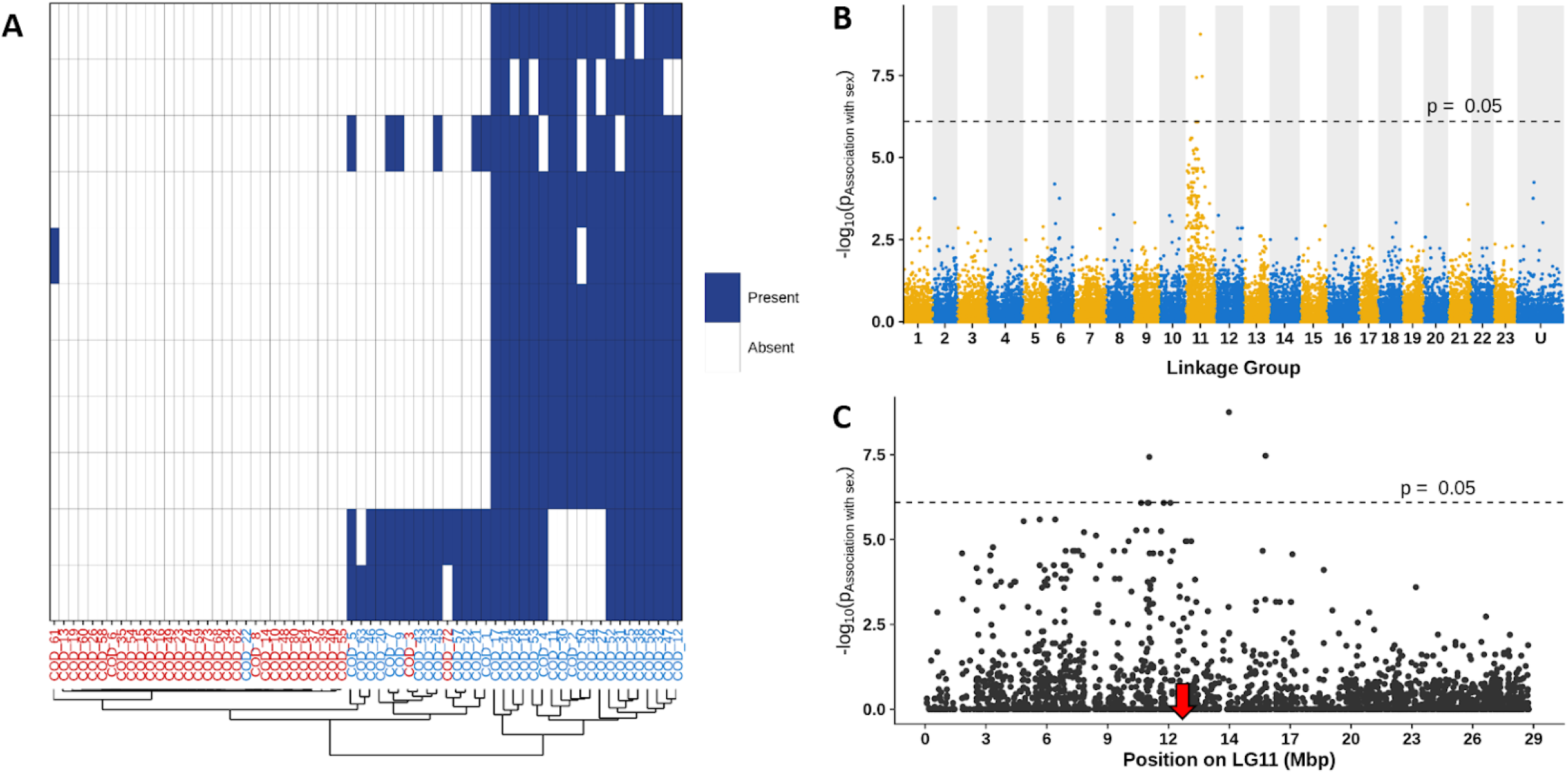
Results of RADSex analysis on *Gadus morhua*. **A**: Tile plot showing the presence of markers in each individual for all markers present with depth higher than five in at least 20 males and in fewer than two females. Each row corresponds to a marker and each column corresponds to an individual, with females in red and males in blue. **B:** Manhattan plot showing −log10(P) of a chi-squared test on the number of males and number of females for all RADSex markers aligned to the genome of *G. morhua*, with odd linkage groups (LGs) in yellow with light background and even LGs in blue with shaded background. Unplaced scaffolds were combined in a super scaffold (U) by decreasing order of size. The dashed line indicates a p-value of 0.05 after Bonferroni correction. **C**: −log10(P) of a chi-squared test on the number of males and the number of females for all RADSex markers aligned to LG11 of *G. morhua*. The dashed line indicates a p-value of 0.05 after Bonferroni correction. Markers significantly associated with sex (p < 0.05) and other markers strongly associated with sex were aligned to the first 20 Mb of the chromosome. The position of the candidate master sex determining gene is indicating by a red arrow.

In the nine other species (**Table 1** and **Supp. Fig. 3**), we did not find any marker significantly associated with phenotypic sex, which indicates that these species either 1) have a small undifferentiated sex locus, 2) lack a genetic sex determination mechanism, or 3) rely on a complex combination of environmental and genetic sex determination factors that prevented the detection of a clear sex-biased signal in the dataset.

## DISCUSSION

We demonstrate that RADSex is an efficient and, thanks to its simple usage and visualization options, user-friendly tool to identify sex-determination systems and sex-biased markers from RAD-Seq data. Unlike other popular RAD-Seq analysis software, *e.g.* STACKS (J. M. Catchen et al., 2011) and PyRad (Eaton, 2014), which catalog polymorphic loci, RADSex creates monomorphic markers. RADSex therefore counts each allele at a locus defined as polymorphic by other software as a separate marker. Working with monomorphic markers allows statistical comparison of populations using straightforward presence/absence tests, yielding results that are comparable between datasets. In contrast, the outcome of grouping sequences into polymorphic markers can be sensitive to parameter values (Paris et al., 2017; Rodríguez-Ezpeleta et al., 2016; Shafer et al., 2018). Leveraging this property of monomorphic markers, RADSex offers a simple and reproducible way to identify *de novo* genetic differences between females and males and to locate these differences on a genome when a reference assembly is available. In addition to this simplicity, RADSex requires few resources and provides comprehensive visualization tools to assist the interpretation of its output.

In particular, the tile plot visualization provides a comprehensive overview of the RAD-Seq results across all individuals and allows the user to quickly identify straightforward cases of genetic sex determination systems. For instance, XX/XY or ZZ/ZW monofactorial systems can be easily detected in absence of any major environmental effects on sex, as demonstrated by our analyses on the ayu (*P. altivelis*) and the tench (*T. tinca*) datasets: in both species, the tile plot revealed a complete sex-linkage of several markers significantly associated with male phenotype, indicating a male heterogametic sex determination system. Besides these straightforward cases, the tile plot visualization is most useful to identify the sex-determination system in more complex datasets, for instance in the presence of outliers and sequencing biases. These complex datasets can then be further explored with the marker depths heatmap visualization, for instance to confirm whether sex-biased markers are absent or present in specific individuals. Using this heatmap, we were able to quickly identify sex-reversed individuals responsible for the non-complete sex-linkage of sex-biased markers in a publicly available medaka dataset (Wilson et al., 2014), as well as outliers in our own common carp (*C. carpio*), Atlantic cod (*G. morhua*), banded knifefish (*G. carapo*), and common molly (*P. sphenops*) datasets. These outliers could arise from human errors in assessing phenotypic sex; in particular, in the common molly, males are externally indistinguishable from females until puberty, which occurs late in some males, and immature testes resemble non-reproductively active ovaries. Because in this species sex was determined from secondary sex characters and macroscopy of the gonads, but not histology, it is possible that the relatively high number of females in which we found male-biased markers were immature males that were mis-sexed as females. However, the presence of outliers could also hint at a more complex sex determination system, for instance polygenic sex-determination (Moore & Roberts, 2013), autosomal modifiers that are polymorphic in the population (Wu, 1983), or simple monofactorial genetic sex determination systems with some sex-reversed individuals triggered by environmental factors (Baroiller & D’Cotta, 2016; Dupoué et al., 2019; Wessels et al., 2017). With a high occurrence of outliers, sex-biased markers may no longer be significantly associated with phenotypic sex, yet these markers can still be biologically relevant.

Because RADSex creates non-polymorphic markers, the number and distribution of markers between individuals is only affected by a single parameter controlling the minimum depth to consider a marker present in an individual. An overly high minimum depth value can lead to false negatives, *i.e.* markers considered absent from an individual because of insufficient sequencing depth, and a very low value can create false positives, *i.e.* markers considered present in an individual because of sequencing errors. Moreover, because the probability of association with phenotypic sex for a marker is adjusted using Bonferroni correction, the stringency of the test for significance increases with the total number of markers identified in the dataset, which is affected by the minimum depth parameter. In the 15 datasets that we generated, median sequencing coverage in an individual was between 30x and 100x, except for three species for which it was lower than 30x (**Supp. Fig. 2**). For all species except the three species with a low sequencing coverage, we found that minimum depth values between 5 and 10 provided a good balance between stringency and minimizing false negatives. However, some datasets may require using low minimum depth values, for instance the three datasets for which median individual sequencing depth was low, or species with large genome sizes like some sharks (Hara et al., 2018) or some amphibians (Sclavi & Herrick, 2019) for which sequencing all individuals of a population with sufficient depth can be costly. In such cases, the memory efficiency of RADSex enables the analysis of large datasets with minimum depth as low as one, which can require a lot of memory with other software.

The numbers of markers significantly associated with phenotypic sex can provide an estimate of the size of the sex determining region, with a low or high number of sex-specific markers reflecting a small or a large non-recombining region, respectively. It is worth noting, however, that other factors can affect this number, for instance the level of differentiation between males and females within the non-recombining region, or bias in GC content within this region (Sigeman et al., 2018; Smeds et al., 2015) potentially leading to an over- or under-representation of RAD-Seq markers depending on the restriction enzyme used for library construction. An extreme case is a complete absence of markers associated with phenotypic sex for any value of minimum depth, which we observed in nine of our datasets. These null results may be linked to two key features of sex determination in teleosts. First, early cytological studies have shown that the majority of teleost fish carry homomorphic sex chromosome pairs (Devlin & Nagahama, 2002), and more recent genomic analyses have revealed that small sex loci are common in teleosts, with the extreme case of a single SNP being the sole difference between the sexes in Fugu (Kamiya et al., 2012). In such cases when there is very little differentiation between the sex chromosomes, the fragmented resolution of RAD-Seq means that the the likelihood to obtain RAD sequences from the sex locus can be very low. Although this problem might be overcome by increasing the number of RAD-seq reads, for example by using a restriction enzyme that cuts more frequently, genome-wide approaches like pooled sequencing of males and females (Feron et al., 2020; Gammerdinger, Conte, Baroiller, D’Cotta, & Kocher, 2016; Wen et al., 2019) or individual whole genome sequencing (Star et al., 2016) may be necessary to identify the sex-determining region. Second, a complete absence of sex-biased markers can reflect a complex sex determination system, for instance a polygenic system or a system involving a strong environmental effect that would weaken the association between genetic markers and phenotypic sex. In fish, many environmental components can interfere with genetic sex determination mechanisms (Kikuchi and Hamakuchi 2013, Heule et al., 2014). Although the visualization tools included in RADSex facilitate the identification of sex reversed individuals, the association between phenotypic sex and genomic regions may still be difficult to detect when sex-reversal is frequent, especially when associated with phenotyping uncertainty. For such systems, pooled sequencing may not be well-suited because male and female pools will each contain heterogeneous genotypes, and individual sequencing remains expensive despite decreasing sequencing costs. A hybrid strategy using both RAD-Seq and pooled sequencing may prove the most efficient for these complex sex-determination systems, as demonstrated in the case of the goldfish (Wen et al., 2019), a species with a strong thermal effect on sex determination (Goto-Kazeto et al., 2006). Finally, in some species sex is determined entirely by environmental factors and therefore does not involve any genetic mechanism (Martínez-Juárez & Moreno-Mendoza, 2019).

One of the strengths of RAD-Seq is the ability to compare populations without a reference genome, which may not always be available for non-model species. However, when a genome is available, locating sex-biased markers provides information on the size and level of differentiation of the sex-determining region and can help to identify candidate genes involved in sex determination. The manhattan plot, circos plot, and contig plot visualizations included in RADSex quickly display the location of the sex-biased markers over the whole genome or over selected regions to identify the sex-determining region. Furthermore, the sex-bias metric and probability of association with sex computed with RADSex can be more effective than F_ST_ computed on polymorphic loci to detect variability in the sex-determining region. This was illustrated by our reanalysis of a publicly available Japanese medaka dataset (Wilson et al., 2014), in which we detected multiple Y chromosome haplotypes that were not found in the original study. Intra-specific sex chromosome polymorphisms have been reported in different taxa, including mammals in which Y chromosome polymorphisms are widely used for human evolution studies (Jobling & Tyler-Smith, 2003) and some fruit flies in which the Y chromosome polymorphism is thought to control male fitness (Chippindale & Rice, 2001). In fish, sex chromosome polymorphisms have been reported in different populations of the guppy, *Poecilia reticulata*, and in other closely related species of guppies (Indrajit Nanda et al., 2014), and different sex chromosomes have even been found within the same species, for instance in tilapias (Cnaani et al., 2008). However, to the best of our knowledge, there are no reports of intra-specific population differences of a single sex chromosome in fish, and our results provide the first example of such a Y chromosome polymorphism in this clade.

As part of this study, we used RADSex to investigate the sex determination systems of 15 species sampled broadly across the ray-finned fish phylogeny (Sup. figure X). We identified XX/XY SD systems in six of these 15 species, and we did not find evidence of a ZZ/ZW SD system in any of the other nine species. Although our sampling is too limited to make general inferences about teleost sex determination, the predominance of XX/XY over ZZ/ZW SD systems in our datasets is in agreement with previous findings that in teleosts, transitions from ZZ/ZW to XX/XY SD systems are more frequent than the reverse (Pennell, Mank, & Peichel, 2018), which would result in XY SD systems being prevalent in this clade.

All our datasets were generated using single-digest RAD-Seq, but the workflow accepts double-digest RAD-Seq data as well. Furthermore, although all species included in this study are fish, we did not use any fish-specific assumptions in RADSex’s implementation, and therefore the computational workflow can be applied to RAD-Seq data from any species. Finally, we specifically developed RADSex to study sex determination, and this design decision is reflected in the wording of this manuscript. However, the computational workflow was designed to be generic, and therefore both the *radsex* software and the *sgtr* R package could be used for other large and major quantitative trait loci with contrasting binary phenotypes.

## ACKNOWLEDGEMENTS

We thank all colleagues who helped us improve the beta version of RADSex by testing it and providing feedback. This project was supported by funds from the “Agence Nationale de la Recherche”, the “Deutsche Forschungsgemeinschaft” (ANR/DFG, PhyloSex project, 2014-2016, SCHA 408/10-1, MS), the National Institutes of Health (USA) grant R01GM085318, JHP), and the National Science Foundation (USA) Office of Polar Programs (grants PLR-1247510 and PLR-1444167 to HWD and grant OPP-1543383 to JHP, TD, and HWD). The MGX core sequencing facility was supported by France Genomique National infrastructure, funded as part of “Investissement d’avenir” program managed by Agence Nationale pour la Recherche (contract ANR-10-INBS-09). RF was partially supported by Swiss National Science Foundation grant PP00P3_170664 to RMW. Common carp were provided by the PEARL INRA 1036 U3E experimental facilities that are supported by the ANAEE-France National Infrastructure. We thank Allyse Ferrara, Quenton Fontenot, and the Bayousphere Lab at Nicholls State University (Thibodeaux, LA) for support with spotted gar sample collection and preparation, Dan Rosauer and Jonathan Meerbeek from the Iowa Department of Natural Resources for walleye sampling. The authors thank the captain and crew of the ARSV *Laurence M. Gould* and the personnel of the US Antarctic Support Contractor for assistance in Chile, at sea, at Palmer Station, Antarctica, and logistically in Denver, CO. Part of this work was supported by a contribution from the Marine Science Center at Northeastern University.

## DATA ACCESSIBILITY

All RAD-Sequencing experiments have been submitted to Genbank under the bioproject PRJNA548074. A computational workflow implementing all the analyses performed in this study, including generating figures, is available at https://github.com/RomainFeron/paper-sexdetermination-radsex. RADSex is released under GPLv3 license; source code, installation instructions, and documentation are available at https://github.com/SexGenomicsToolkit/radsexex. *sgtr* is released under GPLv3 license and available at https://github.com/SexGenomicsToolkit/sgtr.

## AUTHOR CONTRIBUTIONS

RF designed and implemented RADSex and sgtr, with feedback from QP, MW, and YG. YG, JHP, and MS designed the PhyloSex project and advised on results interpretation. QP, MW, BI, JA, AH, KK, ASR, KD, SK, CK, JHP, MS and YG participated in the analysis of the results. HP and LJ prepared libraries and performed the sequencing. RF, QP, JHP, MS and YG drafted the manuscript. RF, QP, RMW, JHP, MS, and YG revised the manuscript. YG, MS, JHP, EJ, SK, MW, MA, CW, BM, AA, TD, FWG, MK, HWD, MO, RN, TS, MN, AW, ØK and IB collected, sexed and/or extracted and prepared gDNA samples. All authors approved the final manuscript.

## SUPPLEMENTARY MATERIAL

**Supplementary Figure 1:**
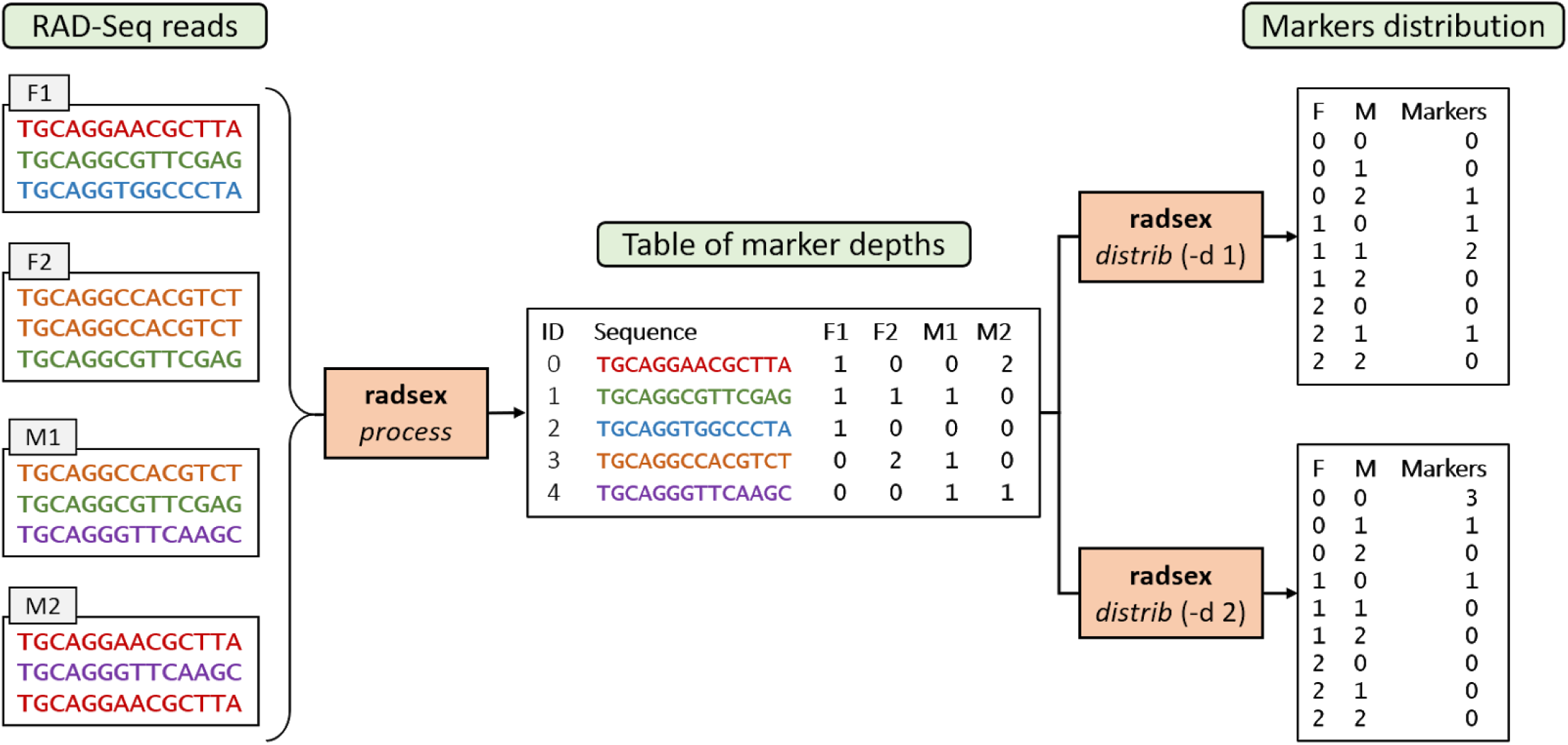
example illustrating the computation of a markers depth table with *radsex process* and the distribution of markers between sexes with *radsex distrib*. RAD-Seq read depth in each individual is summarized in a table of marker depths with *the process* command from *radsex*. This table is used by the *distrib* command to compute the distribution of markers between males and females for a given minimum depth value to consider a marker present in an individual (-d).

**Supplementary Figure 2:**
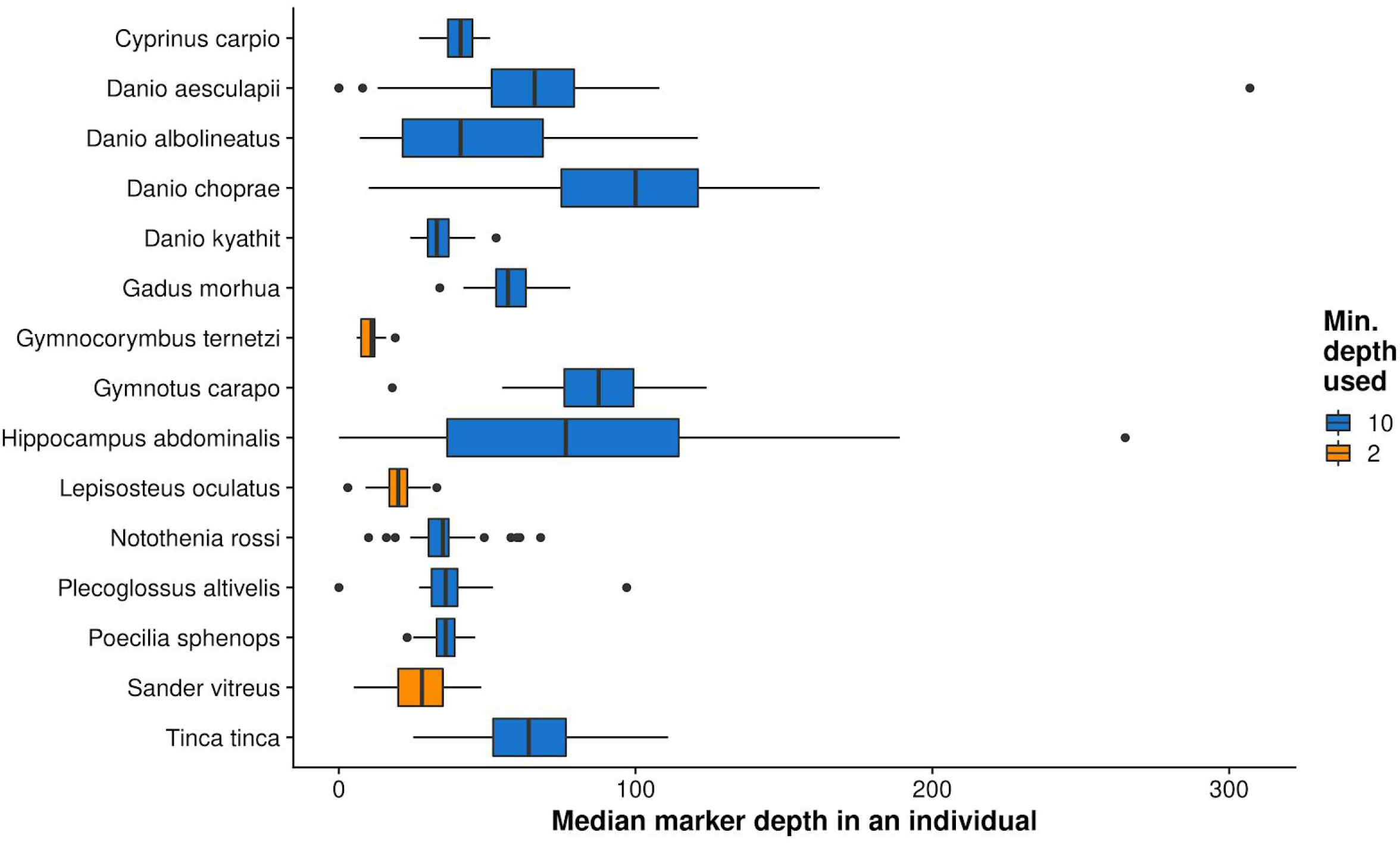
Median marker depth in an individual for each dataset, computed with *radsex depth*. Boxplots show the median, first and third quartiles, and lower and upper extremes of the distribution, with outliers represented as points. Median depth was low for *G. ternetzi*, *L. oculatus*, and *S. vitreus*, and therefore analyses were performed with a lower minimum depth (-d 2 instead of 10) to consider a marker present in an individual.

**Supplementary Figure 3:**
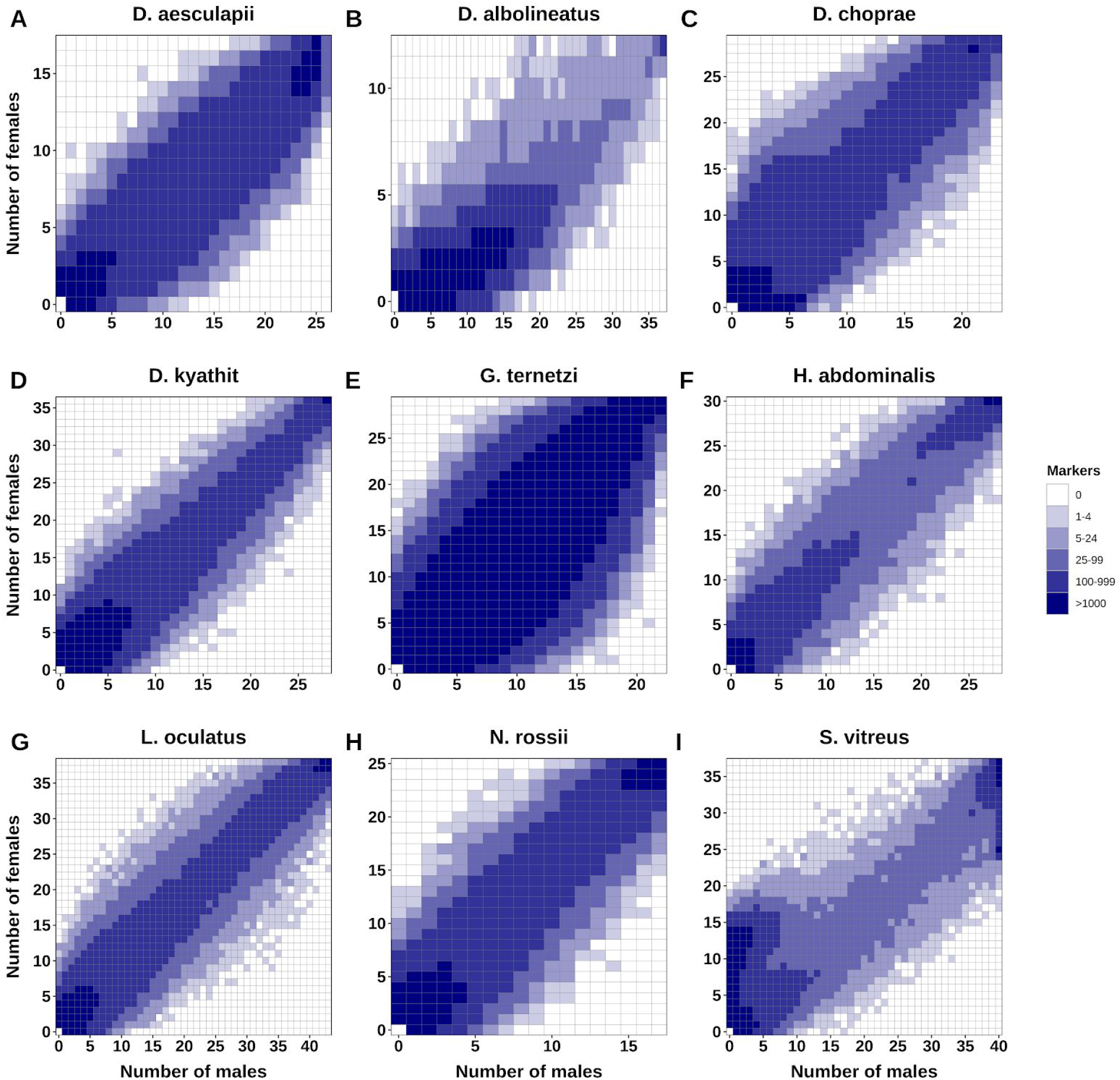
Distribution of markers between males and females in datasets for which no marker significantly associated with sex was found: *D. aesculapii* **(A),** *D. albolineatus* **(B),** *D. choprae* **(C),** *D. kyathit* **(D),** *G. ternerzi* **(E),** *H. abdominalis* **(F),** *L. oculatus* **(G), N** *. rossii* **(H), and** *S. vitreus* **(I)**. The number of markers, given by a tile’s color intensity, is plotted for every combination of number of males (horizontal axis) and number of females (vertical axis). Distributions were computed with a minimum depth of 10 (-d 10), except for *G. ternerzi*, *L. oculatus* and *S. vitreus* for which distributions were computed with a minimum depth of 2 (-d 2).

**Supplementary Table 1:** example of *process* output for three individuals and three markers. In order: marker ID, marker sequence, and depth of this marker in each individual. For instance, the marker with ID 1 corresponds to the sequence TGCAGG..CT (the sequence was shortened), which is present 17 times in the sequencing results of Individual_1, 13 times in individual_2, and 15 times in Individual_3.

| id | sequence | Individual_1 | Individual_2 | Individual_3 |
| --- | --- | --- | --- | --- |
| 0 | TGCAGG..AT | 0 | 15 | 12 |
| 1 | TGCAGG..CT | 17 | 13 | 15 |
| 2 | TGCAGG..GA | 14 | 0 | 18 |

**Supplementary Table 2:** example of *distrib* output for two males and two females. In order: number of males, number of females, number of markers present in the associated numbers of males and females, p-value of association with sex and significativity of this p-value for these numbers of males and females. For instance, there are 950 markers found in 1 male and 2 females in this dataset. This 1 male and these 2 females are not necessarily the same individuals for all 950 markers.

| Males | Females | Markers | P | Signif |
| --- | --- | --- | --- | --- |
| 0 | 1 | 7261 | 1 | False |
| 0 | 2 | 2089 | 0.507763 | False |
| 1 | 0 | 2768 | 0.948911 | False |
| 1 | 1 | 1065 | 1 | False |
| 1 | 2 | 950 | 1 | False |
| 2 | 0 | 715 | 0.424083 | False |
| 2 | 1 | 673 | 0.925099 | False |
| 2 | 2 | 520 | 1 | False |

**Supplementary Table 3:** example of *map* output for four markers. In order: marker ID, contig and position of the contig where the marker was aligned, sex bias, p-value of association with sex and significativity of this p-value for each marker.

| Marker | Contig | Position | Bias | P | Signif |
| --- | --- | --- | --- | --- | --- |
| 0 | Chr1 | 241356 | 0.0689655 | 0.422281 | False |
| 1 | Chr5 | 17595862 | -0.0851293 | 0.604401 | False |
| 2 | Chr7 | 136598 | 0.931035 | 1.69831e-12 | True |
| 3 | Scaffold1 | 365231 | 0.0150862 | 1 | False |
| 4 | Chr13 | 5397682 | 0.0592672 | 0.671211 | False |

**Supplementary Table 4:**
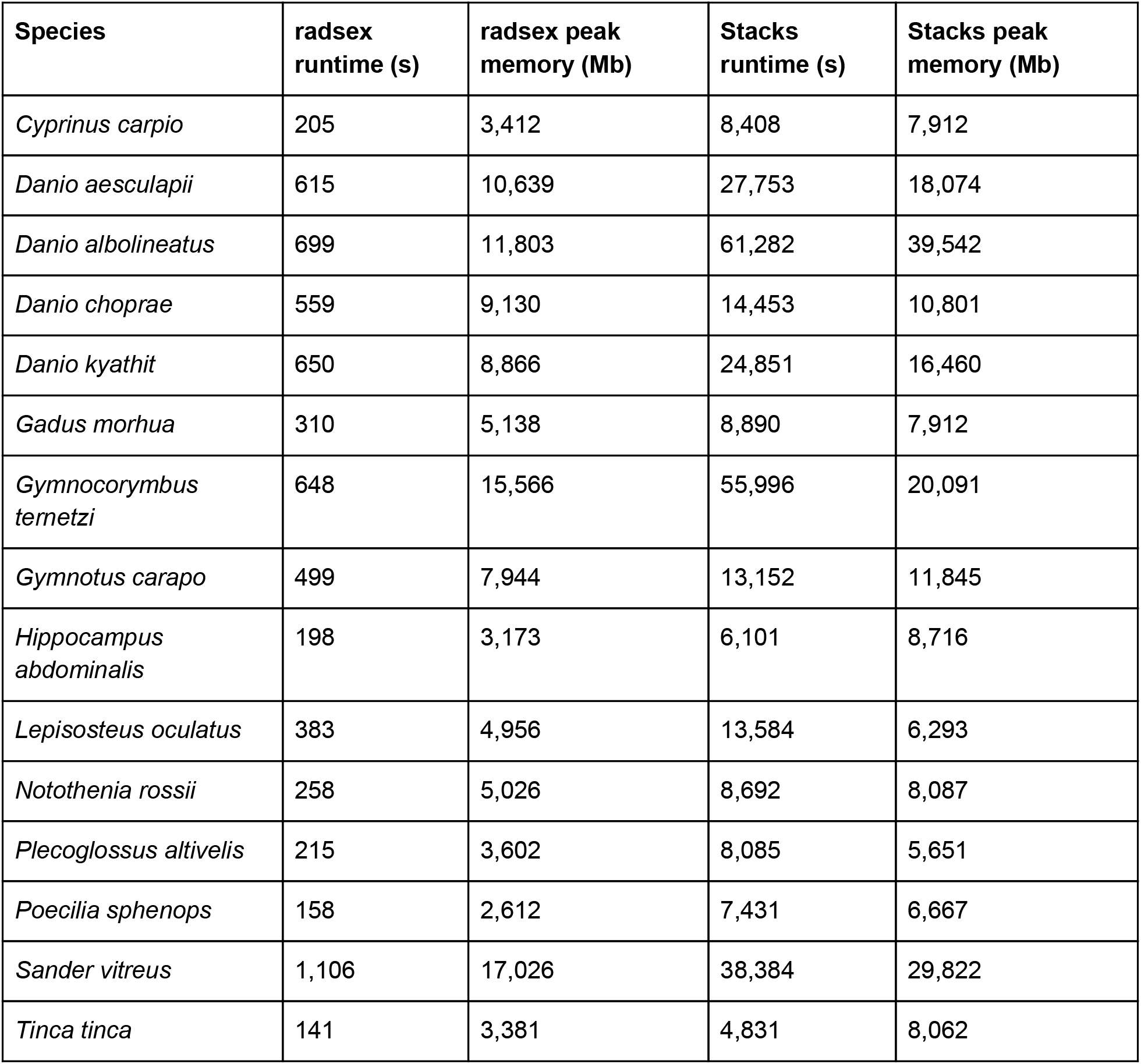
runtime (in seconds) and peak memory usage (in Mb) of *radsex* and *Stacks* for all 15 datasets.

**Supplementary Table 5:** Excel file containing information on sample collection.

**Supplementary Table 6:**
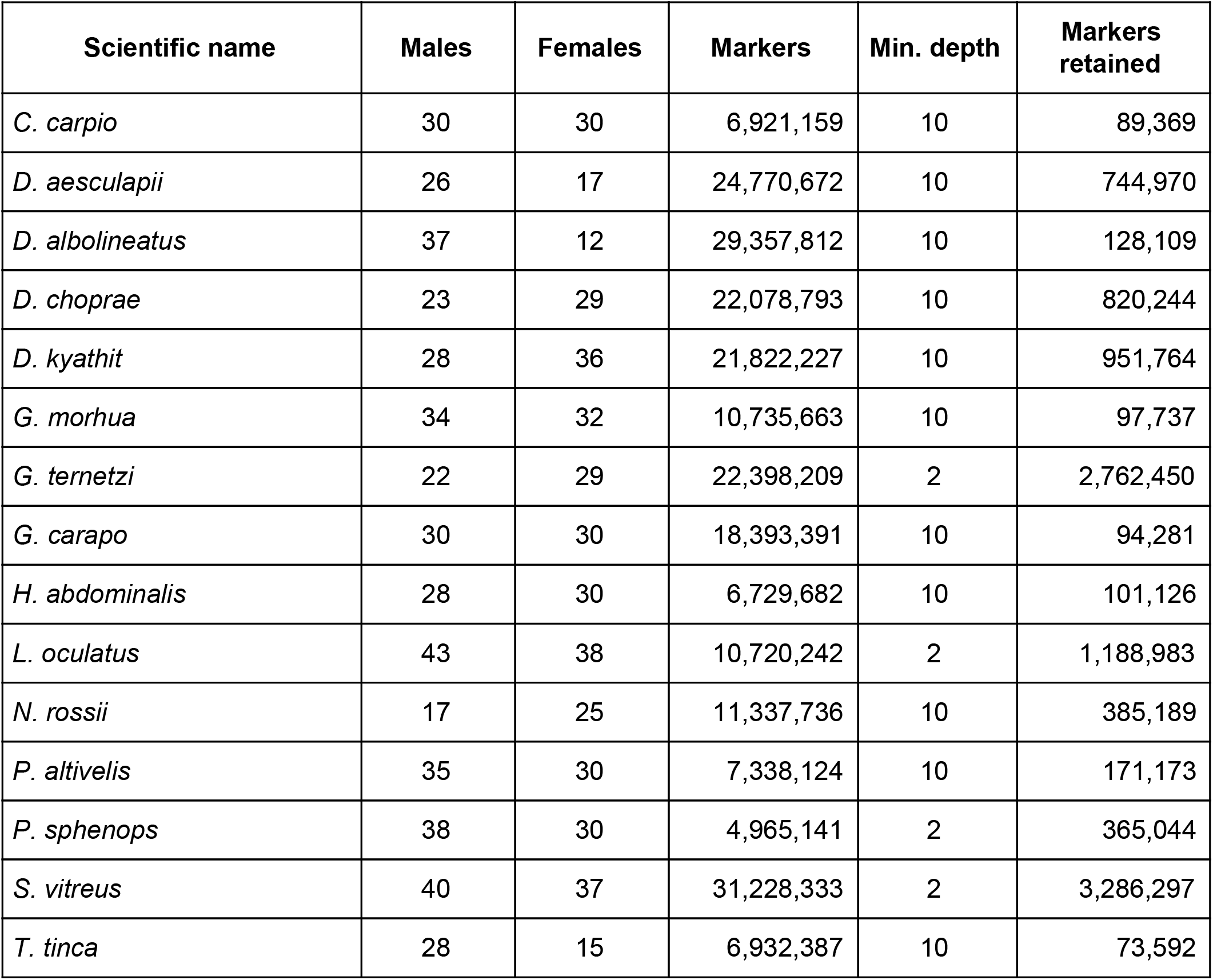
Excel file containing all significant sex-biased marker sequences and the distribution of these markers in all the animals analyzed in *Cyprinus carpio* (7 markers, m = 10), Gadus morhua (3 markers, m = 10), Gymnotus carapo (8 markers, m = 10), Plecoglossus altivelis (4 markers, m = 10), Poecilia sphenops (7 markers, m = 2) and Tinca tinca (6 markers, m = 10).

**Supplementary Table 7:** number of phenotypic males and females, minimum depth to consider a marker present in *radsex*, and number of markers for each dataset. Markers retained correspond to the number of markers present with depth higher than Min. depth in at least one individual.

## Notes

https://github.com/RomainFeron/paper-sexdetermination-radsex

### Competing Interest Statement

The authors have declared no competing interest.

